# Optimal marker gene selection for cell type discrimination in single cell analyses

**DOI:** 10.1101/599654

**Authors:** Bianca Dumitrascu, Soledad Villar, Dustin G. Mixon, Barbara E. Engelhardt

## Abstract

Single-cell technologies characterize complex cell populations across multiple data modalities at unprecedented scale and resolution. Multi-omic data for single cell gene expression, *in situ* hybridization, or single cell chromatin states are increasingly available across diverse tissue types. When isolating specific cell types from a sample of disassociated cells or performing *in situ* sequencing in collections of heterogeneous cells, one challenging task is to select a small set of informative markers to identify and differentiate specific cell types or cell states as precisely as possible. Given single cell RNA-seq data and a set of cellular labels to discriminate, scGene-Fit selects gene transcript markers that jointly optimize cell label recovery using label-aware compressive classification methods, resulting in a substantially more robust and less redundant set of markers than existing methods. When applied to a data set given a hierarchy of cell type labels, the markers found by our method enable the recovery of the label hierarchy through a computationally efficient and principled optimization.

---

Single cell RNA-seq (scRNA-seq) has generated a wealth of data allowing researchers to measure and quantify RNA levels in single cells at unprecedented scales [Macosko et al., 2015, Zheng et al., 2017]. These studies yield valuable insights regarding intrinsic properties of cell type, which is critical to understanding cell development and disease [Zhu et al., 2018]. When coupled with other data modalities, such as measurements of cell surface protein levels [Stoeckius et al., 2017] or spatial transcriptomics [Codeluppi et al., 2018], a more precise and complex partition of the cell type landscape emerges.

Three single cell methodologies motivate our work. First, single cell RNA-sequencing (scRNA-seq) performs short read RNA-sequencing on disassociated cells in a sample; a key goal of scRNA-seq analysis is to label each of the cells in the sample with a precise cell type by considering the genes that are expressed in the cell. Second, spatial transcriptomic experiments use single-molecule fluorescence in situ hybridization (smFISH) approaches [Raj et al., 2008]. These techniques rely on fluorescent probes that bind near genes of interest that are to be quantified called *markers*. When the marker genes bound by probes are expressed above a threshold, fluorescence can be detected using microscopy at the location of expression. Similarly, sorting techniques such as FACS sorting rely on designing a small number of probes for markers that can accurately distinguish cell types according to differences in expression of a small set of cell surface markers [Veluchamy et al., 2017].

If we are to use these methodologies to visualize, characterize, and distinguish cell types among collections of heterogeneous cells, a key challenge from these emerging technologies is in labeling each assayed cell with a precise cell type or cell state label. To do this, gene panels must be designed that allow us to distinguish between cell types both efficiently (with as few markers as possible) and at high precision (discriminating similar cell types). The number of markers is experimentally constrained by the product of the number of fluorescence channels and the number of hybridization cycles [Codeluppi et al., 2018]. In particular, state-of-the-art methods use on the order of 40 markers. The task of optimally choosing markers among all genes that most reliably and precisely distinguish the cell type labels given a hierarchy partitioning the cell types is a combinatorially difficult problem.

Existing approaches to marker selection are scarce and only allow for the identification of markers that distinguish a single cell type from all of the other cell types in a sample [Finak et al., 2015, McDavid et al., 2012]. These methods identify markers that are differentially expressed across two groups by comparing within-group expression with across-group expression. These approaches ignore both hierarchical relationships of cell types and correlations in expression patterns among genes. The simplistic one-versus-all representation of cell types to select markers prevents a solution to the problem when the number of cell types is larger than the number of markers that can be used in an experiment [Reboredo et al., 2016].

In contrast, label hierarchy-aware approaches have the ability to select markers that partition the labels at layers that are not the leaves of the tree, allowing genes that are robustly expressed across a subset of cell types to be selected as markers. For a bifurcating hierarchy the number of markers used is *k*-1, where *k* is the number of cell types (i.e., leaves in the hierarchical tree). Yet, these markers will be redundant if the latent dimension of the space spanned by the gene expression profiles of each cell type is lower than the number of cell types we aim to distinguish. Assuming this is true, then a number of markers smaller than *k* is needed to maintain the hierarchy.

To fill this methodological gap, we developed scGeneFit, a rigorous and efficient approach for marker selection in the context of scRNA-seq data with a given hierarchical partition of labels. Our method draws from ideas in compressive classification [Reboredo et al., 2016] and largest margin nearest neighbor algorithms [Weinberger and Saul, 2009, McWhirter et al., 2018]. scGeneFit shows good performance in both simulated data and in scRNA-seq data. Where traditional approaches test the discriminatory value of each gene separately, our approach jointly recovers the optimal set of genes of a given size that allows robust partitioning of the given labels. Our framework generalizes to settings where the input label partition is captured in a hierarchical, or tree-like, structure.

Briefly, scGeneFit works as follows. Given samples (cells) in a high-dimensional feature space (genes), and corresponding categorical sample labels (e.g., cell type, cell state), label-aware compression methods [McWhirter et al., 2018] find a projection to a low dimensional subspace (space of markers, where the dimension is specified) where samples with different labels remain separated when projected into that space. To ensure that the low-dimensional results enable marker selection, scGeneFit additionally constrains the projection such that each of the subspace dimensions must be aligned with a coordinate axis in the original space. With this constraint, each projection dimension— a single marker—corresponds to a single gene, and not a weighted linear combination of many genes. Fortunately, this constraint eases the computational challenge of the original projection problem, which is intractable in its most general form—turning the optimization into a linear program. For input to our method, we use post-quality-control scRNA-seq data with unique molecular identifier (UMI) counts, a target marker set size, and a hierarchical taxonomy of cell labels. When cell labels do not exist, labels may be inferred using a clustering algorithm, or via another data modality Stoeckius et al. [2017]. Similarly, the label hierarchy for input to our method can be expert-provided Zeisel et al. [2015], or inferred via a hierarchical clustering algorithm.

We assessed scGeneFit’s performance in both simulated scenarios and existing scRNA-seq data. We first investigated the behavior of scGeneFit in the context of simulated data that is often used to evaluate methods for spectral clustering. Samples of dimension *d* +2 are generated such that *d* features are each drawn from a Gaussian distribution with mean zero and variance *σ* = 1, and two features, encoding the desired clusters, represent the two dimensional coordinates on one of two concentric circles with different radii; the cluster label represents whether the sample is on the inner or the outer circle (Fig. 1 A; see Methods). We found that scGeneFit selected as markers the two features representing the circle coordinates whenever the size of the target marker set was ≥ 2, enabling the recovery of the sample label. In contrast, methods that query each of the features independently are not able to identify the two dimensional coordinates.

**Figure 1:**
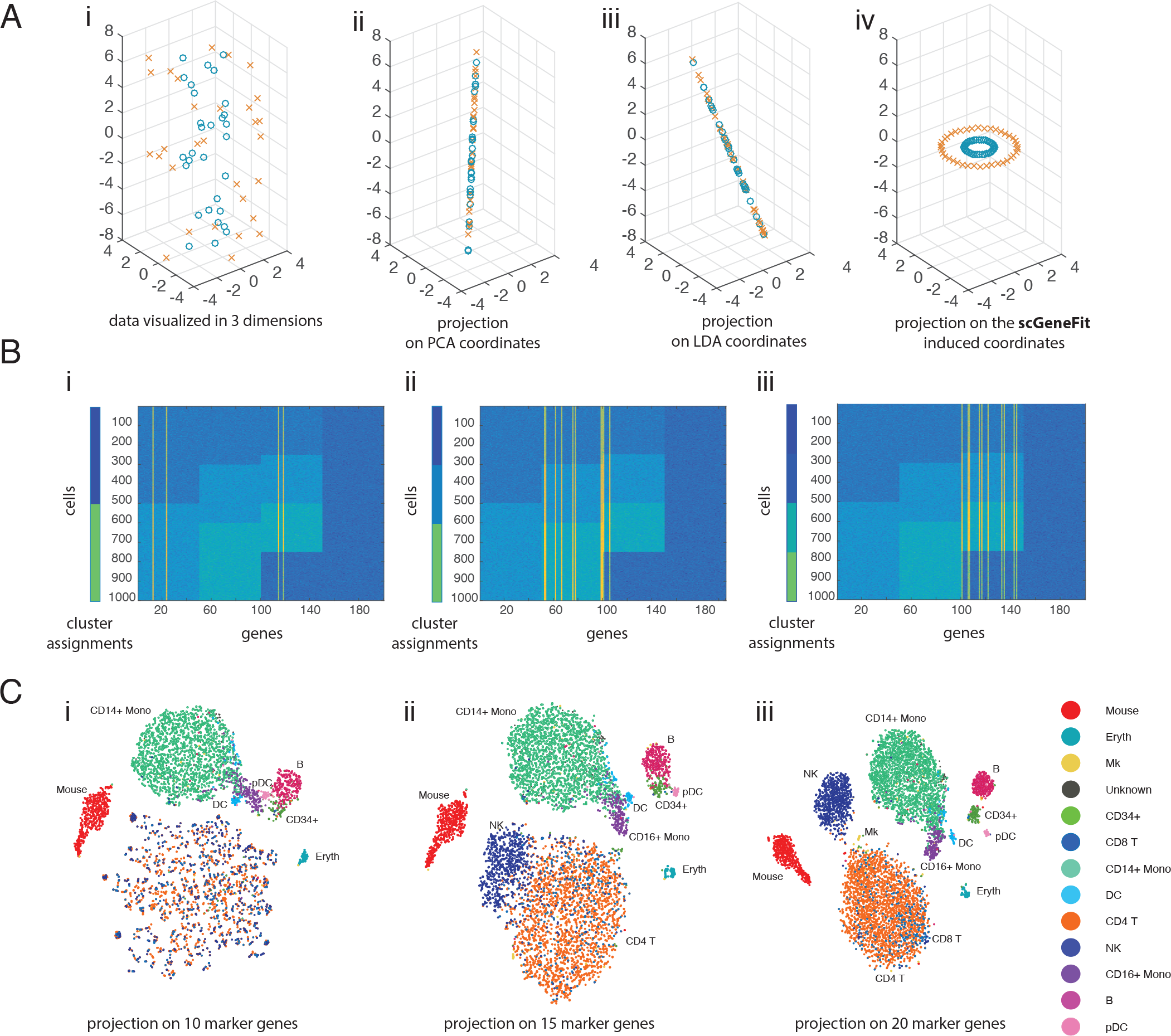
scGeneFit identifies markers associated with a flat partition of cell type labels when applied to simulated data and to scRNA-seq data. **Panel A:** Proof of concept (cf. [McWhirter et al., 2018]); cells are color coded with labels. In simulated high-dimensional data, for each sample, two dimensions (x-and y-axes) are drawn from concentric circles, and the remaining dimensions are drawn from white noise. The underlying structure is not apparent from the data (A-i). Considering each dimension in isolation, marker selection fails to capture the true structure (A-ii,iii). In contrast, scGeneFit recovers the correct dimensions as markers, and is able to recapitulate the label structure (A-iv). **Panel B:** Discriminative markers were correctly recovered by scGeneFit for simulated samples drawn from mixtures of Gaussians corresponding to three label sets with two (B-i), three (B-ii), and four (B-iii) labels, respectively. Each row is a single sample and each column is a single feature, and the shade of blue corresponds to the label assignment, with samples with the same label sharing a color. The yellow lines correspond to the markers selected by scGeneFit. **Panel C:** t-SNE visualization of results from single-cell expression profiles of cord blood mononuclear cells (CBMC) given a flat partition of labels Stoeckius et al. [2017]. scGeneFit reveals that 15 marker genes are sufficient to distinguish 13 distinct cell populations.

Next, we simulated data corresponding to fantasy gene expression profiles from *n* = 1000 cells and *d* = 200 genes (Fig. 1 B). We considered three different labelings of these *n* cells and assigned 50 genes as markers of each partition and another 50 genes as non-markers. Within each of the three labelings, the 50 markers for each label are drawn from a multivariate Gaussian with a 50 dimensional mean and a dense 50 × 50 covariance matrix. The 50 genes that are not markers are drawn from a univariate Gaussian distribution with mean 0 and variance 1. We set the marker set size to 20 and include as input one of the three labelings. We find that scGeneFit recovers markers of the appropriate labeling. Notably, since the third labeling is a finer partition of the first labeling, our method selects markers for both the first and third labelings when given the first labeling, but not vice versa.

We studied the performance of scGeneFit in the context of two scRNA-seq studies. To do this, we applied scGeneFit to a cord blood mononuclear cell (CBMC) study containing isolated cells from cord blood [Stoeckius et al., 2017]. In total, 8, 584 single-cell expression profiles of CBMCs from the top 500 highly variable genes from both human and mouse cell lines were considered. For the cell type hierarchy, we used the reported transcriptome-based clustering that partitions CBMC types into B cells, T cells, natural killer cells (NK), monocytes (CD14+, CD16+), dendritic cells (DC, pDC), erythrocytes, and erythroblasts. We found that the discovered marker sets were able to recover nearly identical cell type partitions in the CBMCs. We explored the space of marker set size close to the number of cell type labels, and see that this number affects the ability to discriminate among cell types (Fig. 1C). We quantified the discriminatory power of the identified marker set by performing *k*-nearest neighbor clustering on the projections of cells in low-dimensional marker space (see Methods). We found that the distinctions between cell type labels are largely preserved in the reduced dimensional space (Fig. 1C). scGene-Fit achieved slightly better performance with fewer markers than the curated marker set based on the differential association (8.05% scGeneFit label classification error versus 9.90% curated marker set error in held-out data; see Methods), and substantially outperformed a random marker set (30% error in held-out data).

Similar performance is achieved in scRNA-seq data characterizing mouse cortical cell type diversity [Zeisel et al., 2015], where we have a expert-derived hierarchy of cell types available. Among the 48 cell types, cells in the mouse somatosensory cortex are organized into a hierarchy governed by four main neuronal types: pyramidal neurons, interneurons, astrocytes, oligodendrocytes, and microglia. We applied the hierarchical version of scGeneFit to identify 30 marker genes that would allow recovery of the hierarchical label structure (Fig. 2A). The selected markers (Supplementary Fig. S1) enabled accurate recovery of the hierarchy of cell types. In particular, the organization of the second level of the recovered hierarchy included two mistakes: the choroid type astrocyte cells tended to be associated with endothelial cells, and the interneurons Int3 were misclassified as pyramidal SS (Fig. 2B). These results coincide with those obtained considering more than 50% more markers in the one-vs-all setup.

**Figure 2:**
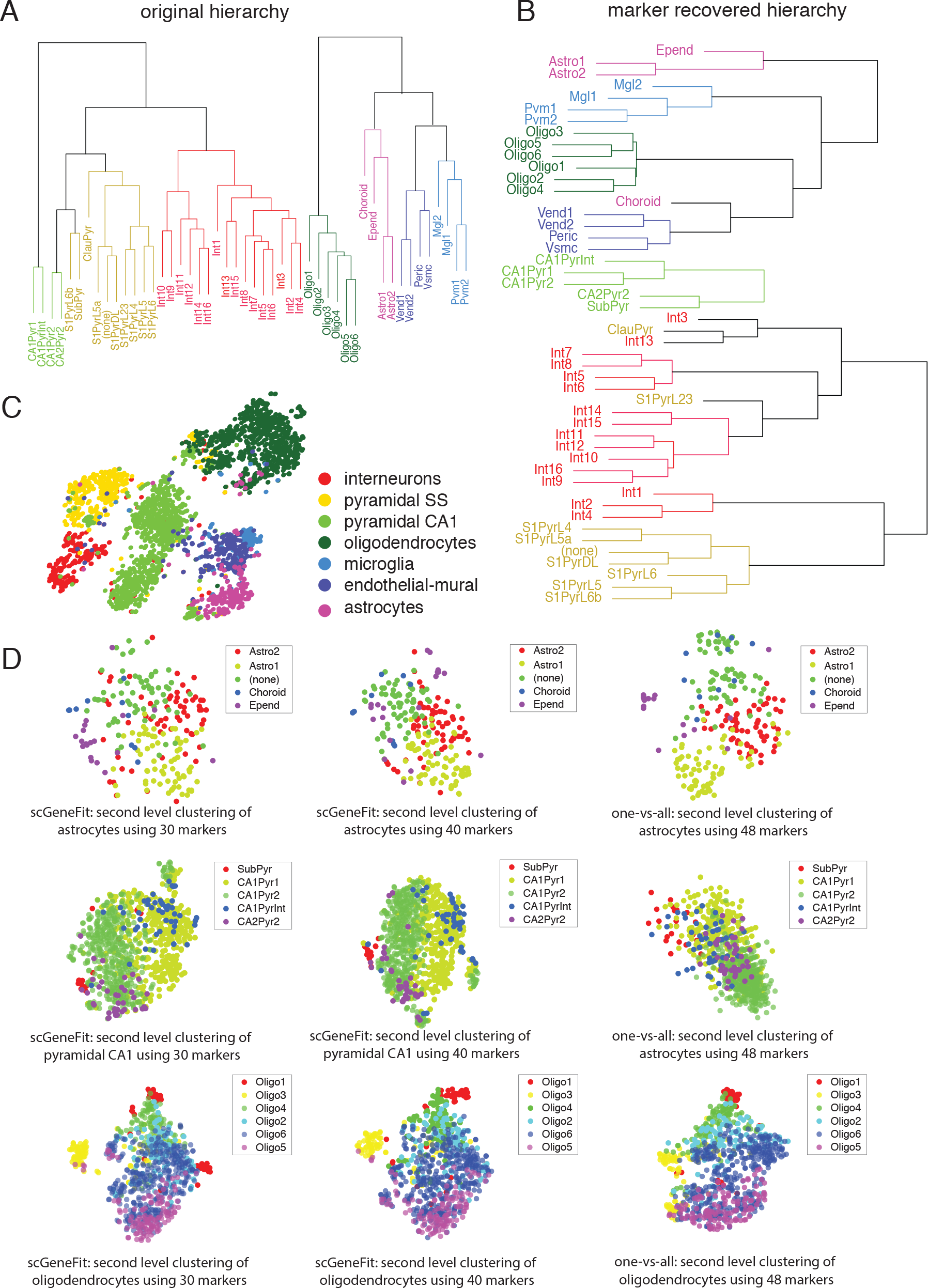
scGeneFit applied to scRNA-seq data with a given hierarchical labeling. **Panel A:** Hierarchical labeling of brain scRNA-seq data Zeisel et al. [2015]. **Panel B:** Hierarchical labeling of the data with respect to the 30 markers chosen by **scGeneFit**. Minimal structural changes are observed when compared to the original hierarchy, using 40% less markers. **Panel C:** a t-SNE plot of the main cell types: interneurons, pyramidal SS cells, oyramidal CA1 cells, oligodendrocytes, microglia, endothelial-mural cells, and astrocytes. **Panel D:** Level two subclustering across three major groups: astrocytes, pyramidal CA1, and oligodendrocytes. t-SNE plots illustrate the performance achieved with scGeneFit with 30 and 40 is comparable with an one-vs-all approach using 48 markers.

We compared the results obtained by scGeneFit at the second layer of the hierarchy using 30 and 40 markers with the one-vs-all method using 48 markers (Fig. 2D). While the subpopulation of microglial cells were poorly distinguished by either method, possibly due to the presence of rare cell subtypes, our method with 30 and 40 markers performed on par with the one-vs-all method with 48 markers (classification error at first layer of hierarchy on held-out cells: 10.21% (30) 6.77% (40) 12.42 (one-vs-all); Fig. 2D). In particular, scGeneFit performs well in the astrocytes, pyramidal CA1, and oligodendrocytes subpopulations (Fig. 2D; Supplementary Table S1).

## Discussion

In summary, we develop a method, scGeneFit, that identifies markers to distinguish cell types given a structured partition (flat partition or hierarchy) of cell type labels for a specific tissue type. We show that scGeneFit is able to accurately recover a set of markers of a pre-specified size that enables robust labeling of cell types in scRNA-seq data. scGeneFit is able to handle more complex relationships among class labels than a flat partition, making it the first approach to exploit this structure to robustly solve the marker selection problem in an efficient and principled fashion. In the future, we envision relaxing the categorical labeling to a manifold constraint that will allow selection of markers to place unlabeled cells at, for example, time points along cell trajectories or locations in spatial assays.

## Methods

### scGeneFit

In the marker selection problem, *x*_*i*_ ∈ ℝ ^*d*^ is the gene expression measurements of the *i*th cell for *d* different genes. We assume the subset of the cells used for training include labels.

#### Setup

We model the marker selection problem as a label-aware dimension reduction method inspired by compressive classification and largest margin nearest neighbor algorithms [McWhirter et al., 2018]. One such method, SqueezeFit, aims to find a projection to the lowest dimensional subspace for which samples with different labels remain farther apart than samples with the same label. Consider a data set 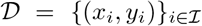 in ℝ^*d*^ × [*k*], here *x*_*i*_ is a sample and *y*_*i*_ is its corresponding label. We denote 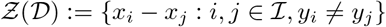 as the vector difference between samples with different labels.

The following optimization problem corresponds to finding the orthogonal projection to the lowest dimensional space that maintains a prescribed separation Δ > 0 between samples with different labels:

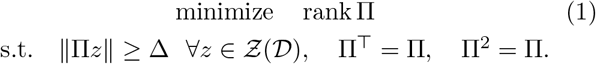

Here, Π is the low-dimensional projection, and Δ > 0 is the desired minimum distance between projected samples Π*x*_*i*_ and Π*x*_*j*_ with different labels. This parameter reflects a fundamental tension in compressive classification: Δ should be large so as to enable sufficient separation of samples with different labels in the low dimensional space, and simultaneously the projected space rank Π should be of low dimension so that this projection effectively reduces the dimension of the sample. To address the intractability of the optimization in (1), a convex relaxation technique is used [McWhirter et al., 2018]:

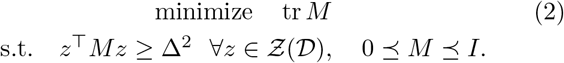

The relaxation extends the feasible set from the set of orthogonal projections where optimization is intractable – matrices Π that satisfy the constraints in (1) – to the set of positive semidefinite matrices – matrices *M* that satisfy the constraints in (2) – where one can use standard optimization toolboxes to find the global optimum in polynomial-time [Grant and Boyd]. The trace norm of *M* corresponds to the *ℓ*_1_-norm of the vector of eigenvalues of *M*. Therefore, minimizing the trace norm tr*M* encourages *M* to be low-rank [Srebro and Shraibman, 2005].

#### Marker selection

scGeneFit finds a prescribed number of gene markers, so that when the samples are projected onto those marker dimensions they exhibit the same separation of cells with different labels as in the original gene space. The objective of selecting a handful of marker genes in mathematical terms translates to finding a projection onto a subset of coordinates; specifically, *M* is diagonal matrix with entries *α*_1_, …,*α*_*d*_. This constraint simplifies the optimization to a linear program:

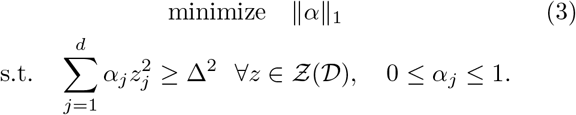

The objective’s same *ℓ*_1_ trace norm promotes sparsity in the matrix *M* [Srebro and Shraibman, 2005]. As a result, numerical experiments show that the solution of (3) is in fact sparse, and the dimension of the projection—the number of selected markers—is smaller than the dimension of the original space.

In order for this method to be useful in practice, we modify the optimization formulation (3) to allow for outliers, and we specify the dimension of the projected space (i.e., the number of markers) *s*, leading to the scGeneFit optimization problem:

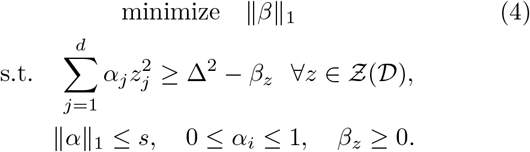

Here *β* is a slack vector that quantifies how much the margin between sets with different labels is violated for each constraint [Cortes and Vapnik, 1995]. *β* is indexed by the elements 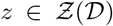 and its dimension equals that of the constraint set 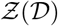.

### Incorporating label hierarchies

Consider a hierarchical partition of the samples denoted by *T_*σ*_* where *σ* is an ordered set of indices. Say *T*_*σ*′_ ⊂ *T*_*σ*_ if *σ* is a prefix of *σ*′ (for instance *T*_*ijk*_ ⊂ *T*_*ij*_ ⊂ *T*_*i*_, corresponding to a three-level hierarchy; see Figure 3 for a concrete example).

**Figure 3:**
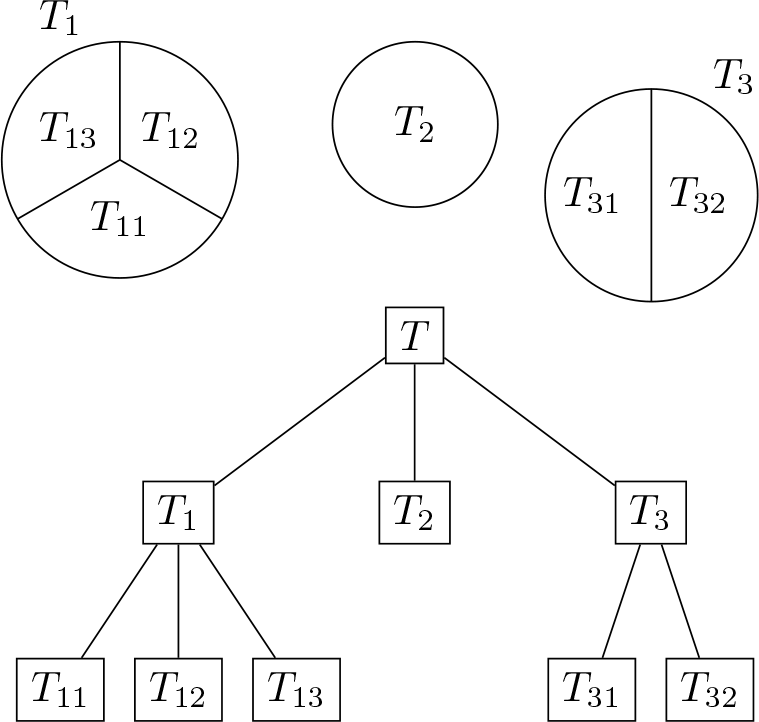
Example of hierarchical partition explaining the notation

When provided with the structured relationship of the labels, scGeneFit solves the optimization problem (4), replacing the set of constraints 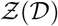 to 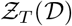 to reflect the hierarchical information. In detail,

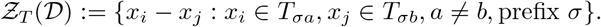

### Alternative optimization constraints

The optimization problem described above for scGeneFit is effective when the label structure is a flat (star shaped) hierarchy; however, when the label structure has additional layers, we would like to add an additional constraint to encourage labels that are closer in hierarchical space to also be closer in the projected (marker) space. In particular, let 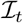 be the set of indices *i* ∈ {1, …, *n*} of cell profiles with label *t*, and let *n*_*t*_ be the number of cells in that set. The projected center of the profiles labeled *t* is Π*c*_*t*_ with 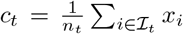.

The desired constraints can thus be formally encoded as 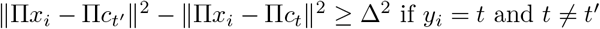.

The hierarchical scGeneFit objective encodes the intuition that the distance between labeled cells should reflect the label distance in the given label hierarchy (Supplementary Information). This is given by the linear program

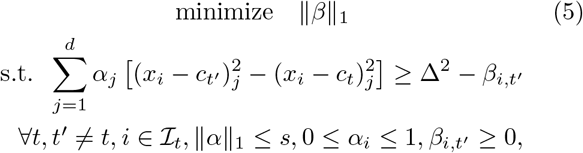

where, as before, *β* is a slack vector.

### Hyperparameter setting

scGeneFit requires specification of two hyperparameters: *s* (the target number of markers) and Δ (the target separation of samples with different labels). In the hierarchical setting, different values of Δ can be chosen for the different levels of the hierarchy. A rule of thumb is to choose Δ as a function of the separation between the classes in ℝ^*d*^.

### Optimization of the linear program and scalability

The optimization problem (4) is a linear program that we solved with Matlab’s built-in solver `linprog`. The computational bottleneck of the linear program is the number of constraints in 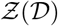, which *a priori* scales quadratically with the number of cells. In order to resolve this issue and make the optimization more efficient, we select the most relevant constraints by considering, for each sample, the *K*-nearest neighbors from each of the other classes. If the number of samples is too large, we randomly select a subsample, run scGeneFit on the subsample, and project the held-out samples using the markers chosen on the subsample.

### Runtime

The computational complexity of linear programming is an open problem in optimization, but it is known to be asymptotically upper bounded by *O*(*N*^2.5^), where *N* is the size of the problem (number of variables plus number of constraints) [Cohen et al., 2018]. For the particular experiments we perform, we solve scGeneFit with 4000 variables and 6000 constraints in less than 40 seconds, on Matlab 2018a running on an Intel Xeon CPU 1.90GHz using less than 4Gb of memory.

### Dataset description and preprocessing

#### Zeisel

Cells in the mouse somatosensory cortex (S1) and hippocampal CA1 region were classified based on 3005 single-cell transcriptomes via scRNA-seq. The nine major molecularly distinct classes of cells (layer 1) were obtained through a divisive biclustering method, and corresponding subclasses of cells (layer 2) were obtained through repeating the biclustering method within each major class [Zeisel et al., 2015].

#### CBMC

The cord blood mononuclear cells (CBMCs) were produced with CITE-seq [Stoeckius et al., 2017]. Single cell RNA data processing and filtering were performed as specified in Stoeckius et al. [2017]. In particular, the data are sparse and normalized by log_2_(1 +*X*).

### Evaluation Metrics

In order to evaluate the performance of scGeneFit we first split the data in training (70%) and test (30%). We train a K-nearest neighbor classifier on the training data (for *K* = 3, 5, 15) after projection to the corresponding markers (computed on the entire dataset). We evaluate the classifier on the test data and report the misclassification error with respect to the known classes (Supplementary Table S1). We also evaluate the performance of *k*-means clustering, using Matlab’s kmeans++ built in implementation, reporting the smallest misclassification error among 10 random initializations. For the hierarchical dataset in Zeisel we evaluate the performance at the second level

### Code availability

The code required to conduct the simulations and reproduce the analyses is available at https://github.com/solevillar/scGeneFit

## References

Simone Codeluppi, Lars E Borm, Amit Zeisel, Gioele La Manno, Josina A van Lunteren, Camilla I Svensson, and Sten Linnarsson. Spatial organization of the so-matosensory cortex revealed by cyclic smFISH. bioRxiv, page 276097, 2018.

Michael B Cohen, Yin Tat Lee, and Zhao Song. Solving linear programs in the current matrix multiplication time. arXiv preprint arXiv:1810.07896, 2018.

Corinna Cortes and Vladimir Vapnik. Support-vector networks. Machine Learning, 20(3):273–297, 1995.

Greg Finak, Andrew McDavid, Masanao Yajima, Jingyuan Deng, Vivian Gersuk, Alex K Shalek, Chloe K Slichter, Hannah W Miller, M Juliana McElrath, Martin Prlic, et al. Mast: a flexible statistical framework for assessing transcriptional changes and characterizing heterogeneity in single-cell RNA sequencing data. Genome Biology 16(1):278, 2015.

Michael Grant and Stephen Boyd. Cvx: Matlab software for disciplined convex programming, version 2.1.

Evan Z Macosko, Anindita Basu, Rahul Satija, James Nemesh, Karthik Shekhar, Melissa Goldman, Itay Tirosh, Allison R Bialas, Nolan Kamitaki, Emily M Martersteck, et al. Highly parallel genome-wide expression profiling of individual cells using nanoliter droplets. Cell, 161(5): 1202–1214, 2015.

Andrew McDavid, Greg Finak, Pratip K Chattopadyay, Maria Dominguez, Laurie Lamoreaux, Steven S Ma, Mario Roederer, and Raphael Gottardo. Data exploration, quality control and testing in single-cell qPCR-based gene expression experiments. Bioinformatics, 29 (4):461–467, 2012.

Culver McWhirter, Dustin G Mixon, and Soledad Villar. Squeezefit: Label-aware dimensionality reduction by semidefinite programming. arXiv preprint arXiv:1812.02768, 2018.

Arjun Raj, Patrick Van Den Bogaard, Scott A Rifkin, Alexander Van Oudenaarden, and Sanjay Tyagi. Imaging individual mRNA molecules using multiple singly labeled probes. Nature Methods, 5(10):877, 2008.

Hugo Reboredo, Francesco Renna, Robert Calderbank, and Miguel RD Rodrigues. Bounds on the number of measurements for reliable compressive classification. IEEE Transactions on Signal Processing, 64(22):5778–5793, 2016.

Nathan Srebro and Adi Shraibman. Rank, trace-norm and max-norm. In International Conference on Computational Learning Theory, pages 545–560. Springer, 2005.

Marlon Stoeckius, Christoph Hafemeister, William Stephen-son, Brian Houck-Loomis, Pratip K Chattopadhyay, Harold Swerdlow, Rahul Satija, and Peter Smibert. Simultaneous epitope and transcriptome measurement in single cells. Nature Methods, 14(9):865, 2017.

John P Veluchamy, María Delso-Vallejo, Nina Kok, Fenna Bohme, Ruth Seggewiss-Bernhardt, Hans J Van Der Vliet, Tanja D De Gruijl, Volker Huppert, and Jan Spanholtz. Standardized and flexible eight colour flow cytometry panels harmonized between different laboratories to study human NK cell phenotype and function. Scientific Reports, 7:43873, 2017.

Kilian Q Weinberger and Lawrence K Saul. Distance metric learning for large margin nearest neighbor classification. Journal of Machine Learning Research, 10(Feb):207–244, 2009.

Amit Zeisel, Ana B Munñoz-Manchado, Simone Codeluppi, Peter Löonnerberg, Gioele La Manno, Anna Juréeus, Sueli Marques, Hermany Munguba, Liqun He, Christer Betsholtz, et al. Cell types in the mouse cortex and hippocampus revealed by single-cell RNA-seq. Science, 347 (6226):1138–1142, 2015.

Grace XY Zheng, Jessica M Terry, Phillip Belgrader, Paul Ryvkin, Zachary W Bent, Ryan Wilson, Solongo B Ziraldo, Tobias D Wheeler, Geoff P McDermott, Junjie Zhu, et al. Massively parallel digital transcriptional profiling of single cells. Nature Communications, 8:14049, 2017.

Lingxue Zhu, Jing Lei, Bernie Devlin, Kathryn Roeder, et al. A unified statistical framework for single cell and bulk RNA sequencing data. The Annals of Applied Statistics, 12(1):609–632, 2018.

