## Supplementary Info for "Optimal marker gene selection for cell type discrimination in single cell analyses"

### SI for scGeneFit : Cluster guided optimal selection of target genes in scRNA-seq

#### Evaluation Metrics

K-nearest neighbor based metrics were used to evaluate the performance of the flat clustering. Due to the small number of points per subcluster this method is not suitable for evaluating the lower level of the hierarchy.

Marker selection for CBMC

| | **original data** | **random markers** | | **one-vs-all** | **scGeneFit** | | |
|---|---|---|---|---|---|---|---|
| markers | 500 | 13 | 20 | 13 | 13 | 20 | 30 |
| $K = 3$ | 7.47 | 19.77 | 14.08 | 9.90 | 9.40 | 8.47 | 8.05 |
| $K = 5$ | 7.20 | 21.35 | 20.41 | 9.01 | 8.82 | 7.97 | 7.93 |
| $K = 15$ | 7.85 | 23.48 | 17.68 | 9.01 | 8.59 | 8.28 | 7.93 |
| $k$-means | 14.99 | 22.45 | 15.55 | 11.60 | 10.92 | 10.70 | 10.84 |

Marker selection for Zeisel (first level cluster)

| | **original data** | **random markers** | | **one-vs-all** | **scGeneFit** | | |
|---|---|---|---|---|---|---|---|
| markers | 4000 | 7 | 40 | 7 | 7 | 30 | 40 |
| $K = 3$ | 2.77 | 64.70 | 28.41 | 12.43 | 16.87 | 10.21 | 6.77 |
| $K = 5$ | 3.44 | 57.82 | 25.75 | 11.87 | 15.65 | 8.54 | 5.88 |
| $K = 15$ | 4.10 | 54.16 | 25.31 | 10.43 | 14.54 | 8.32 | 6.55 |
| $k$-means | 19.78 | 27.28 | 22.17 | 9.74 | 10.86 | 8.75 | 6.44 |

Table 1: Percentage of misclassified CBMC cells (top table) and Zeisel cells (bottom table) from $K$-nearest neighbor and $k$-means after compressing with random markers, markers chosen as most significantly associated with a cluster and scGeneFit (a lower score is a better score). (The Id column takes identity to be the "compression" operator.) For the first rows we split the data in 70% training (6032 cells for CBMC, 2104 for Zeisel) and 30% test (2585 cells for CBMC, 901 for Zeisel). We train a $K$-nearest neighbor classifier on the compressed training set, and report its misclassification error in the compressed test set. The last row considers the smallest misclassification error of $k$-means clustering among 10 runs of $k$-means with different random seeds. Due to the high variance in the number of points per cluster, $k$-means is not a suitable clustering method for this CBMC.

### Detected Markers

scGeneFit identifies both cell-type specific markers (e.g., Pde1a, Cnr1, Enpp2, Mef2c), and markers that show broad patterns of behavior (e.g. the low expression of Tmsb4x is particular to astrocytes). Further interpreting the grammar governing this behavior is subject of future work, and is outside the scope of the current paper.

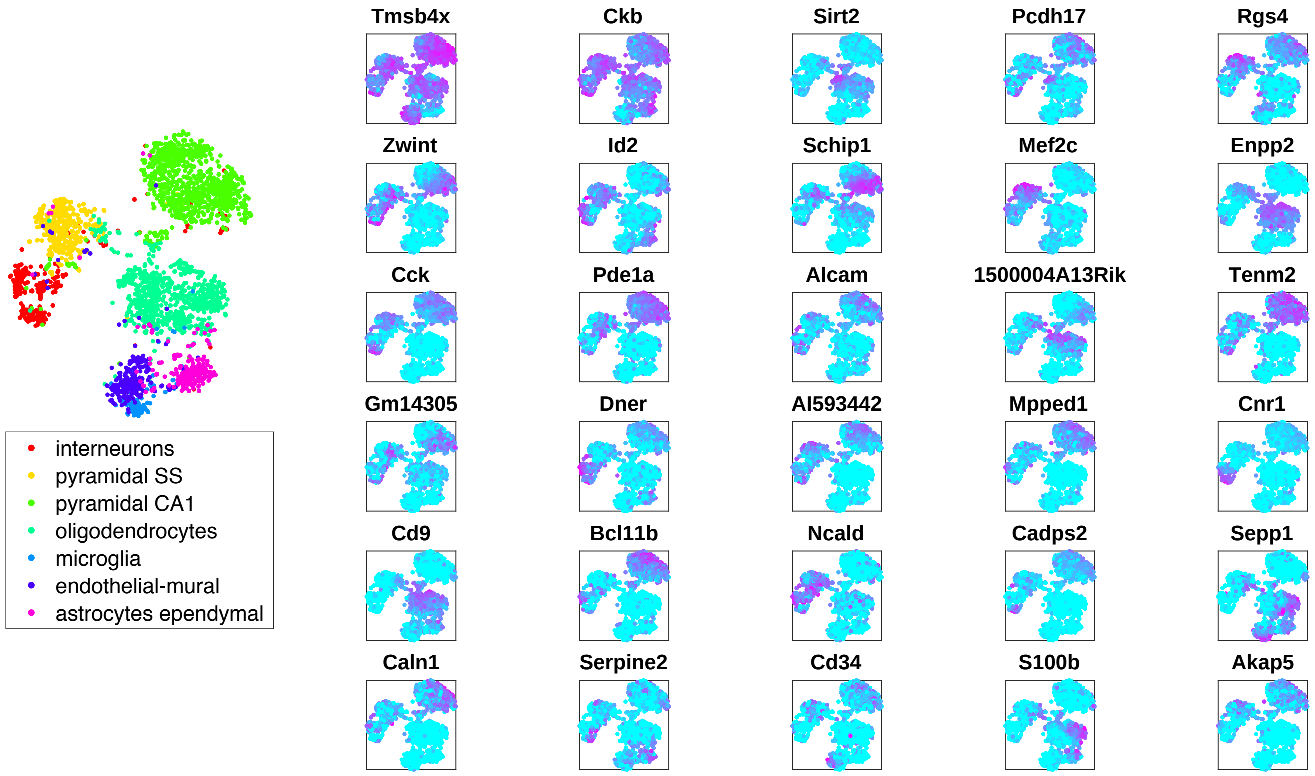


Figure 1: Reference t-SNE plot of the main cell types in the Zeisel data set: interneurons, pyramidal SS cells, oyramidal CA1 cells, oligodendrocytes, microglia, endothelial-mural cells, and astrocytes (left). The 30 markers uncovered by scGeneFit in the hierarchical setting. The markers are overlayed based on their expression – low (blue) to high (pink) – on the reference Zeisel t-SNE plot (right).
